# Gaussian-distributed codon frequencies of genomes

**DOI:** 10.1101/480152

**Authors:** Bohdan B. Khomtchouk, Wolfgang Nonner

**Affiliations:** Department of Biology, Stanford University, 371 Serra Mall, Stanford, CA 94305, USA; Department of Physiology and Biophysics, University of Miami Miller School of Medicine, Miami, FL 33136, USA

**Keywords:** evolution, mathematical modeling, molecular biology, cell biology

## Abstract

DNA encodes protein primary structure using 64 different codons to specify 20 different amino acids and a stop signal. Frequencies of codon occurrence when ordered in descending sequence provide a global characterization of a genome’s preference (bias) for using the different codons of the redundant genetic code. Whereas frequency/rank relations have been described by empirical relations, here we propose a statistical model in which two different forms of codon usage co-exist in a genome. We investigate whether such a model can account for the range of codon usages observed in a large set of genomes from different taxa. The differences in frequency/rank relations across these genomes can be expressed in a single parameter, the proportion of the two codon compartments. One compartment uses different codons with weak bias according to a Gaussian distribution of frequency, the other uses different codons with strong bias. In prokaryotic genomes both compartments appear to be present in a wide range of proportions, whereas in eukaryotic genomes the compartment with Gaussian distribution tends to dominate. Codon frequencies that are Gaussian-distributed suggest that many evolutionary conditions are involved in shaping weakly-biased codon usage, whereas strong bias in codon usage suggests dominance of few evolutionary conditions.

## 1. Introduction

Genome sequencing has produced a large amount of information on how specific genomes use the 64 different codons of the redundant genetic code. This information on codon usage by genomes has raised many issues ranging from fundamental to technical: Can codon usage help elucidate the origin of the universal genetic code [1]? Do genes carry information beyond amino acid sequence in their usage of synonymous codons [2]? What are the evolutionary mechanisms and possible (dis)advantages that make genomes use certain different codons more (less) often than others [2,3,4,5]? How do variations of codon usage arise in evolution [6,7,8,9]? What needs to be understood and controlled to optimize translation in heterologous expression systems [10]?

The detailed information compiled on diversity of codon usages within and across specific genomes has naturally overshadowed a search for common patterns in codon usage that overarch specific codon usages in specific organisms. One such pattern is the descending sequence of genomic codon frequencies. It characterizes bias inherent to the usages of different codons made within a given genome. Intra-genomic bias is necessary for the existence of inter-genomic diversity in the usage of the different codons.

Interest into frequency/rank relations of genomes first concerned linguistic analogies of the ‘language of genes’ and spoken languages [11]. Empirical formal descriptions from this line of work include a power of rank (Zipf’s law) [11,12], an exponential of rank [13,14], and a combination of exponential and linear relations [15]. Codon frequency/rank curves were described by statistical relations with either no external parameter [16], or two external parameters [17,18]. With exception of the statistical model of [16], these descriptions have not been interpreted with regard to characteristics of evolutionary conditions that shape codon usage. In contrast, to account for observations of inhomogeneous codon usages among genes of the same organism, Gusein-Zade and Borodovsky [19] have proposed that there exist two compartments of genes that make distinct usages of the different codons.

In this study we investigate intra-genomic bias in codon usage and test the applicability of a two-compartment model in a large set of genome-wide codon usages. We find that superposition of two patterns of codon frequencies describes genomic frequency/rank plots consistently. Variation of the proportion of the two patterns captures the variations observed among frequency/rank plots of genomes from many taxa. The frequency distributions underlying the two patterns are distinct. In one compartment, prevailing in eukaryotes, codon usage follows a random pattern as described by a Gaussian distribution of frequencies. Usage bias in that compartment is weak. In the other compartment usage bias is strong to the extent of using essentially a subset of the available different codons. Prokaryote genomes reveal both compartments in widely varying proportions.

The existence of a genomic compartment using codons with Gaussian-distributed frequencies likely implies the existence of many contributing evolutionary conditions that together have shaped codon usage in eukaryotic genomes.

## 2. Biased usage of codons: observations

Different codons occur with a range of frequencies in a given genome, and a given different codon tends to occur with generally different frequencies across different genomes. These two aspects of bias in codon usage are illustrated in the cases of four organisms sampled from diverse taxa (Fig. 1A,B).

**Figure 1:**
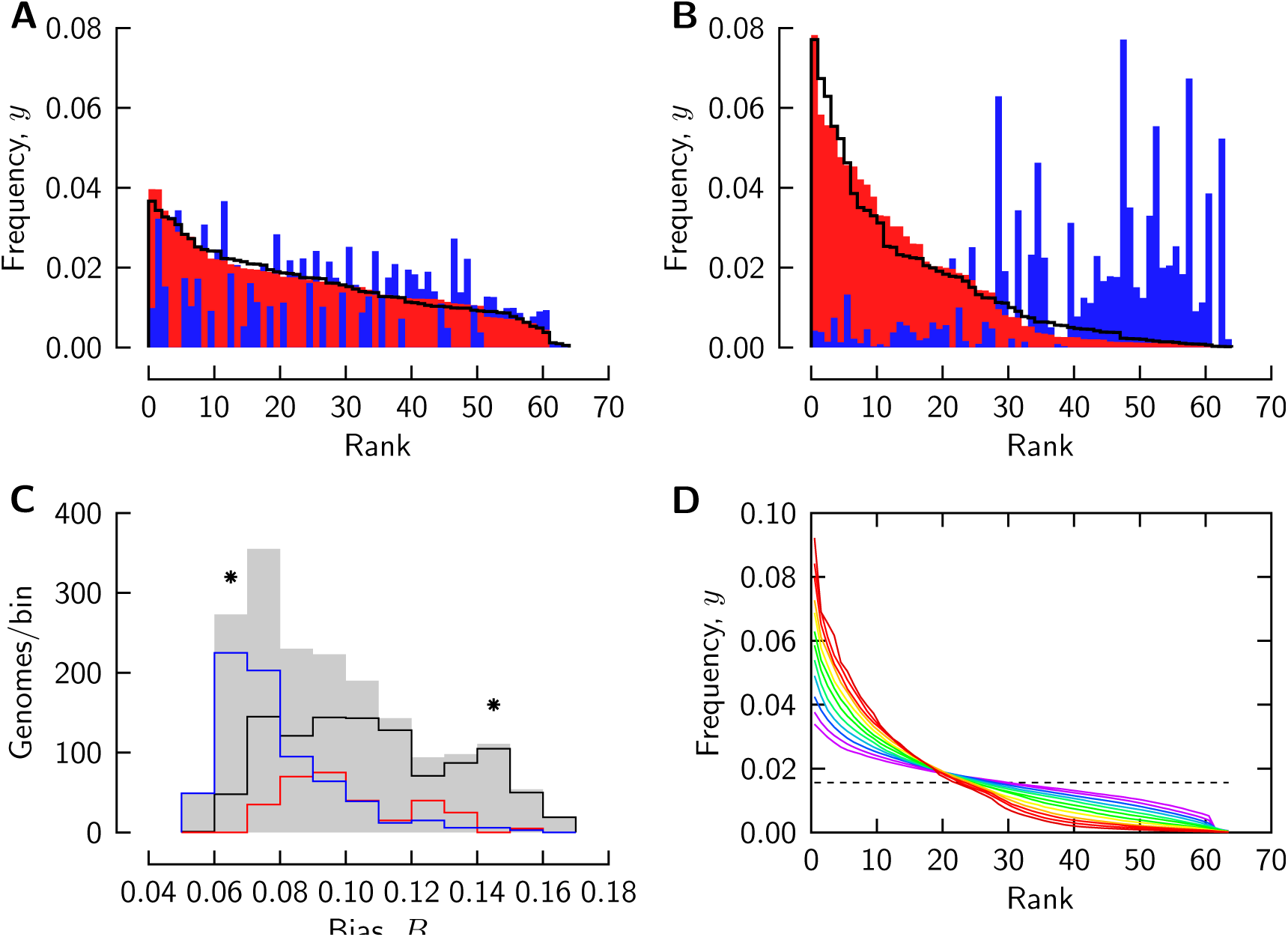
Biased usage of different codons in genomes. Codon frequencies of *Homo sapiens* (A, red color), *Arabidopsis thaliana* (A, blue color), *Streptomyces griseus* (B, red color), and *Clostridium tetani E88* (B, blue color). Frequencies are ordered as described in the text, with the smaller frequency of each two compared genomes shown in the foreground. C: histograms of bias *B* for 1840 genomes from the CUTG database (gray shaded) and of the eubacterial (*black line*), archaeal (*red*, scaled by 5), and eukaryotic subsets (*blue*) of these genomes. The total set comprised 1855 genomes, of which 1840 located to the binned range; the latter are analyzed in this study. Asterisks mark the bins to which the genomes of the panels A,B locate. D: frequency/rank curves averaged over the genomes of each histogram bin in panel C (with bias increasing from purple to red).

In Fig. 1A, the frequencies of the 64 different codons in human genomic DNA have been put in descending order (*red color*). When the frequencies of the different codons of the plant *Arabidopsis thaliana* are plotted in the order of the ordered human codons, they do not form a descending sequence (*blue color*). The codon frequencies of the plant, however, when ordered among themselves, form a descending sequence that is quite similar to that of the ordered human frequencies (*line*). Codon usages of the bacteria *Clostridium tetani* and *Streptomyces griseus* also reveal similar intra-genomic bias, but diverse patterns of cross-genomic bias (Fig. 1B). Both forms of bias are stronger in the bacteria than in human and plant. In each of the pairs of genomes of Fig. 1A,B, a shared pattern of intra-genomic bias is associated with two distinct patterns of cross-genomic bias.

Here we study the intra-genomic aspect of codon usage bias in the genomic coding DNA of a set of 1840 organisms of different taxa tabulated in the CUTG database [20] (files ‘gbxxx.spsum.txt’ where xxx = bct, inv, mam, pln, pri, rod, vrt). We restrict analysis to genomes represented with total codon counts 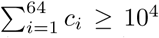, and define frequencies of occurrence, *y*_*i*_, of different codons *i* (1 ≤ *i* ≤ 64) by

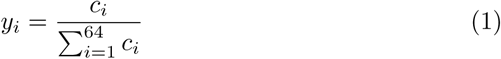

For a non-parametric assessment we measure intra-genomic ‘bias’ as

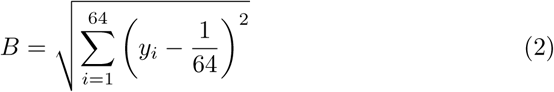

For a hypothetical genome in which all different codons occur with the uniform frequency 1/64, bias *B* = 0. A real genome will have bias *B>* 0.

A histogram of codon usage bias B constructed over the entire set of 1840 analysed genomes is shown in Fig. 1C (*gray-shaded*). Histograms drawn as *lines* represent the subsets eubacteria (*black*), archaea (*red*, counts scaled by the factor 5), and eukaryotes (*blue*). The genomes presented as examples in Fig. 1A,B locate to the bins marked by *asterisks*.

Weak bias of codon usage is characteristic of eukarytotic genomes, whereas prokaryotic genomes reveal a wide range of bias. Overall, bias varies about threefold over the sampled set of genomes.

Frequency/rank curves averaged over the genomes in each bin of the bias histogram in Fig. 1C are shown in Fig. 1D. Codon usage bias with respect to average codon frequency (*dashed line*) is observed in all genomes, but a characteristic of the frequency/rank curve changes in the succession of curves from weak (*purple lines*) to strong bias (*red lines*): in curves associated with weak bias, curvature changes direction near the mean frequency, whereas curves associated with strong bias decline without inflection. Moreover, frequency of genomes with strong bias decays in a cascade-like fashion about the lower and upper halfs of the ranks.

Frequency/rank relations of a genome assign to a given frequency the number of different codons that have frequencies larger than or equal to that frequency. In this way they define the cumulative distribution of the random variable, codon frequency. For this statistical view of the descending frequency sequence, rotate Fig. 1 clockwise by 90 degrees. Then the horizontal axis in panels A,B, and D is the random variable, codon frequency. A position on the vertical axis, read as offsetwith respect to rank 64, indicates how many different codon values have frequencies less than that at that frequency. The distributions in Fig. 1A reveal a sigmoidal cumulative distribution, whereas those in Fig. 1B ressemble an exponential distribution.

In an earlier study of codon usage, Gusein-Zade and Borodovsky [19] have investigated the possibility that genomes are inhomogeneous due to the existence of gene compartments that make distinct usages of different codons. Specifically, they developed a model of two compartments, each characterized by an exponential distribution. When only one compartment was present in a genome, the distribution was exponential. When both compartments co-existed, the overall frequency distribution was described by the convolution of the two scaled exponential distributions. This model was compared to a small data set then available to the authors and found to account for characteristics such as the inflection observed in certain frequency/rank relations.

In view of the larger data set provided by the CUTG database, we noticed that the GZB model with two exponentially distributed components falls short regarding the range of bias that it can describe. Here, bias can vary between 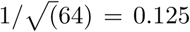 and 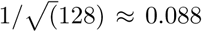 depending on the proportion of the two compartments. This range is substantially less than the observed bias range (Fig. 1C). In this study, we find that a two-compartment model of codon usage describes also the larger data set of the CUTG database. Then variations among genomes are accounted for by a single parameter, the proportion of the two compartments. An improvement of the GZB model, however, is necessary to describe the larger data set: the distributions that describe codon usage in each compartment need to be chosen differently.

In our improved model, one compartment uses different codons according to a Gaussian distribution that produces minimal bias. We suppose this compartment to dominate codon usages like those by the genomes in Fig. 1A. A Gaussian distribution would be expected to arise due to the Central Limit Theorem of statistics if many different evolutionary conditions shape codon usage in this compartment. Alternatively, the Gaussian compartment could represent many smaller compartments with not necessarily Gaussian distributions of codon frequencies.

The other compartment of our model is governed by a distribution that produces even stronger bias than the exponential distributions proposed by GZB. The exponential distribution, as pointed out by GZB [19], ‘obeys the principle of maximum diversity of frequencies’, but this does not exclude that more biased distributions are possible (though less likely to arise by pure chance). For instance, genomes might restrict their machinery of translation to using a subset of different codons within the degenerate genetic code, or even use certain amino acids with preference. Such a codon usage is manifest in the genomes of Fig. 1B. In our model, we derive an empirical description of codon frequencies in the second compartment from a subset of the data. We suppose this compartment to dominate in genomes that use different codons with strong bias.

## 3. Construction of the model

We describe the model together with a Monte-Carlo numerical approach that we use to construct it (Fig. 2). Our approach reveals (and allows one to assess) a limitation inherent to modeling multiple compartments of codon usage.

**Figure 2:**
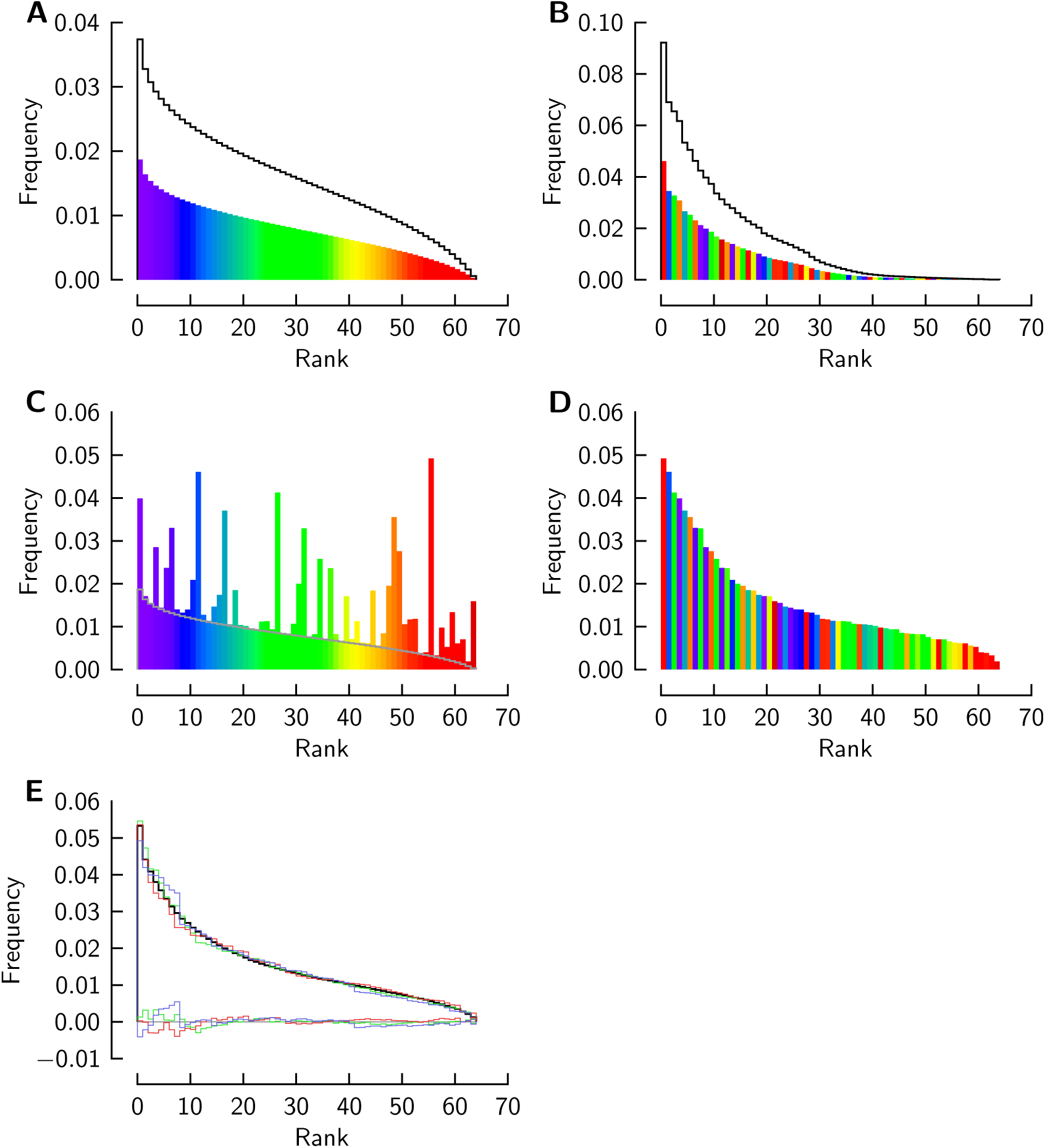
Codon usage in a two-compartment genome model. Frequency sequences computed with Gaussian (A) or empirical frequency distribution (B) (*lines*). *Colored* : frequencies scaled by 1/2 for constructing a joint sequence representing two codon compartments of equal sizes. The colors illustrate different codons associated with the frequency ranks: the order of the second compartment is randomized with regard to the first. (C) Pairwise algebraic sum of the compartmental frequencies of each different codon (matched by color). (D) Frequency sequence (C) after re-ordering for a descending joint sequence. (E) Three samples of frequency sequence like that in (D) computed for different randomizations of the order of different codon in panel B (*colored lines*), and the mean of 100 samples (*black line*). Colored lines along the bottom of the graph: differences between individual samples and the mean.

Assume a genome comprises two compartments in which, generally, each different codon is used according to two distinct and mutually independent statistical distributions: a Gaussian distribution in compartment 1, and an empirically described distribution in compartment 2. Both distributions are naturally truncated to the range of codon frequency, 0 ≤ *y ≤* 1. Overall frequency of a different codon is given by the algebraic sum of the scaled frequencies in the two compartments. If *α* is the proportion of compartment 1,

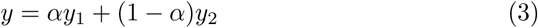

The cumulative probability of frequency *y*_1_ in compartment 1 is that of a truncated Gaussian distribution,

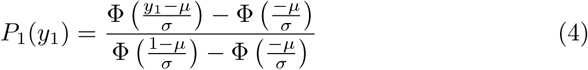

where Φ denotes the cumulative normal distribution function with position *µ* and scale σ. These two parameters are restricted by the requirement that the mean of the truncated distribution be equal to 1*/N*_codons_ (*N*_codons_ = 64 being the number of different codons). This restriction follows from the normalization inherent to codon frequency. It leads to the relation

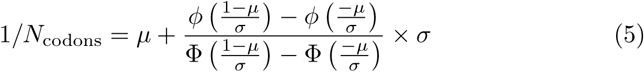

where ϕ (*t*) = *d*<I(*t*)*/dt*. Thereby, the Gaussian distribution has only one external parameter. We will choose *σ* and determine *µ* by solving eq. 5 for *µ*.

We generate a frequency sequence for compartment 1 using Monte-Carlo sampling of frequency into *N*_codons_ discrete bins that are associated with uniform increments of cumulative probability over the range 0 *<P <* 1. In each of 10^6^ sampling cycles, a cumulative probability value is drawn from a generator of uniform random values in the range 0 *< R <* 1. The associated frequency *y*_1_ is computed by solving eq. 4 for *y*_1_ using a root finder. The frequency *y*_1_ is included into the frequency average accumulated in the bin associated with the probability interval. A Monte-Carlo sampled frequency sequence is shown in Fig. 2A (*solid line*). This calculation needs to be done only once for a chosen Gaussian distribution.

We compute the frequency sequence of compartment 2 empirically, as average over a group of genomes that reveal the strongest bias of codon usage in our data set. We average the frequency sequences of the genomes locating to the right-most bin of the bias histogram (Fig. 1C). In this study we will not further analyse the statistical basis of this empirical distribution. We will show here that the characteristics of that distribution are universally detected in genomes across taxa, as are the characteristics of a particular Gaussian distribution.

To construct a frequency sequence for a genome comprising, e.g., two equally large codon compartments, we scale the compartmental distribution frequencies by 1*/*2. We also randomize the association of different codon and frequency rank among the two scaled frequency sequences. This is done by assuming some order of different codons in the ranks of both compartments. Compartment 1 retains that order (represented by a rainbow sequence of colors in Fig. 2A), whereas the order is scrambled in compartment 2 by swapping the colors of randomly chosen pairs of ranks in 10^4^ cycles (Fig. 2B).

We merge the scaled and ordered sequences by algebraically adding to each frequency from compartment 1 the frequency from compartment 2 that is associated with the same different codon (i.e., color). This produces the frequency sequence in Fig. 2C, which happens to be no longer in descending order. Reordering of that sequence produces the joint frequency sequence of the two codon compartments of the genome (Fig. 2D).

The frequency sequence of the compartmentalized genome is intermediate between the compartmental sequences. It also has stochastic roughness that does not exist in the contributing sequences. Both compartmental sequences are thoroughly sampled in the procedure of merging frequencies but each joint frequency is sampled from exactly one pair of frequencies that is formed by a particular different codon in the two compartmental sequences. We cannot expect to model this pseudo-stochastic aspect of joint frequencies because an observed genomic frequency sequence does not reveal how a different codon is associated with rank in each compartment. We can, however, assess the potential consequences of this uncertainty by simulations with the model.

We re-compute the joint frequency sequence using different associations of different codons and frequency ranks in the model compartments (by randomizing the ‘colors’ in compartment 2 for each trial). Fig. 2E shows examples of frequency sequence computed in three different trials (*colored lines*) and an average curve computed over 100 trials (*black line*) after re-ordering the sequence from each trial. All individual trials produce frequency sequences that fluctuate with respect to their mean to an extent small enough (rms of ≈ 10^-3^) to preserve systematic features of the frequency sequence. These simulations quantify how much the mean theoretical curve, which does not capture consequences of specific ranks of different codons, is expected to diverge from the curve observed in a genome in whose compartments different codons have specific, but unknown ranks.

The computer code for all computations was written from scratch in a PostScript-like language and is available under https://github.com/Bohdan-Khomtchouk/Gaussian-distributed-codon-frequencies-of-genomes. Code was executed using a virtual machine [21].

## 4. Comparison of ordered sequences of observed codon frequencies with the theoretical sequences

We test the improved two-compartment model on 1840 genomes from the CUTG database. Frequency sequences of the entire genome set are modeled by adjusting only the proportion *α* of the compartment with the Gaussian distribution while maintaining the Gaussian or empirical distributions of each compartment invariant. The scale of the Gaussian distribution is chosen to be *σ*= 0.009, which implies the position *µ* = 0.1461 by eq. 5. The standard deviation of the truncated Gaussian distribution is 0.0081, and that of the empirical distribution 0.0207. These compartmental distributions (and *α* = 1*/*2) have been used in the computations for Fig. 2. In optimizing *α* we quantify the closeness of fit by the rms residual between observed and mean theoretical frequencies (the latter determined from 100 trials as described above). This residual is expected to approach 10^-3^ for *α* = 0.5 because the theoretical frequency sequence is an average over many different rankings of a different codon’s frequency in the two compartments (Fig. 2E).

Fig. 3A,B,D,E shows observed and theoretical frequency/rank relations for the genomes introduced in Fig. 1A,B. The model describes the data quite well, as is indicated by the residuals of the fit (*dotted lines*).

**Figure 3:**
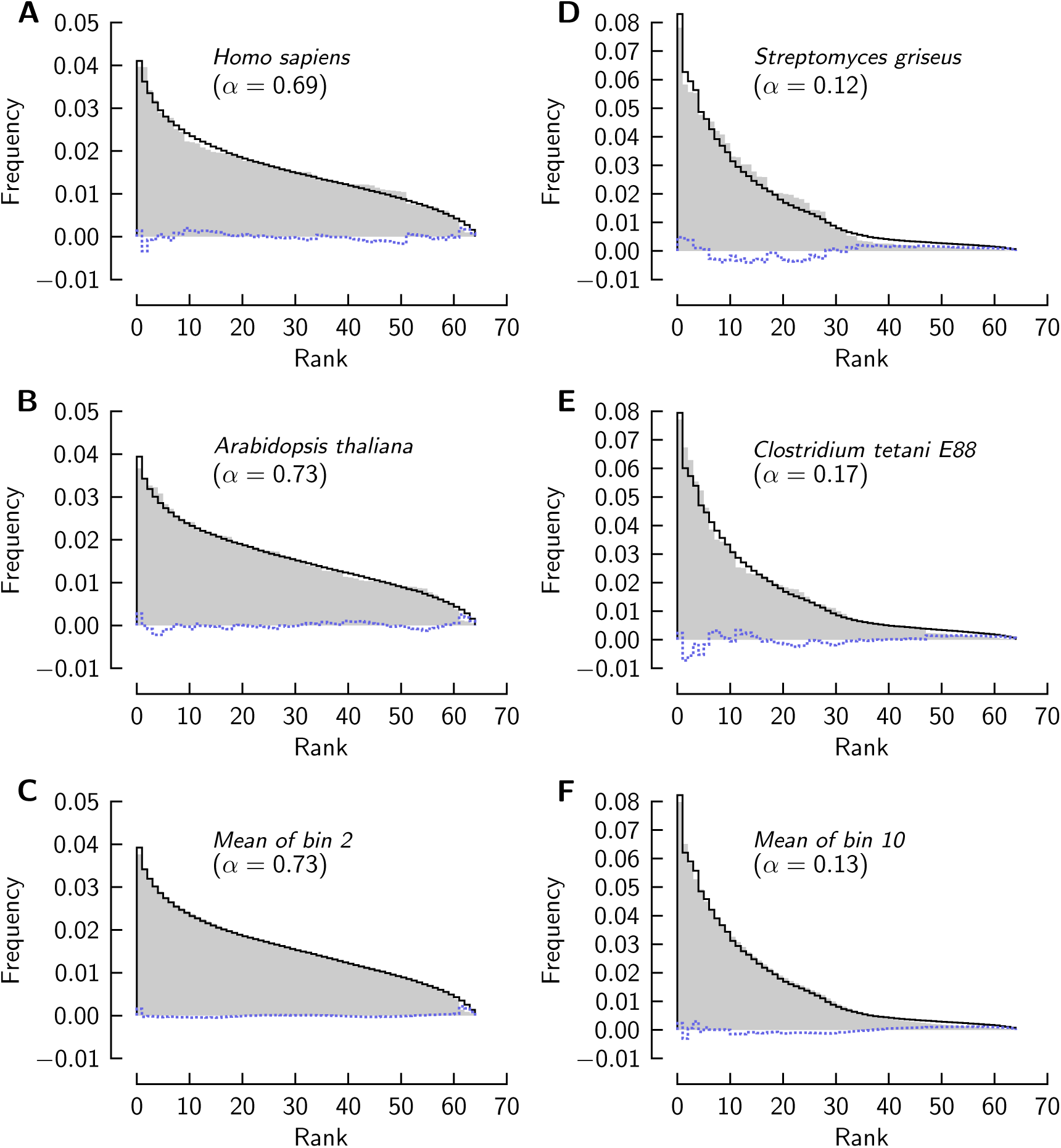
Observed frequencies of codon occurrence compared to theory. Gray shaded: ordered sequences of the frequencies of four genomes (A, B, D, E) and ordered frequencies averaged across the subsets of genomes in the two marked histogram bins in Fig. 1C (C, F). Theoretical frequencies (*black lines*) are computed with the indicated proportion, *α*, of the compartment with Gaussian distribution; *dotted lines*: residuals between observed and theoretical frequencies. The *rms* values of the residuals, scaled by 10^3^, are: 1 (A), 0.8 (B), 0.4 (C), 2.2 (D), 1.9 (E), and 1.1 (E).

The specific order of codons in the hypothesized compartments of the individual genomes, which is unknown, might actually limit our model’s capacity to account for the frequencies in an individual genome. On the assumption that different genomes of similar codon usage bias have different specific orders of codons in their compartments (compare Fig. 1A,B), we expect that averaging the ordered sequences of frequencies of such genomes produces a frequency sequence that is more accurately predicted by the (averaged) sequence of the model than are sequences of individual genomes. Fig. 3C,F compare averaged observed frequency sequence with fitted theoretical sequence. The residuals are substantially smaller than in the case of the individual genomes (Fig. 3A,B,D,E; see legend for rms values of residuals). The model describes frequency/rank relations averaged over genomes more accurately than those of individual genomes of similar usage bias.

Fig. 4 extends the comparison of observed and theoretical frequency sequences to twelve genomes that are commonly studied. These genomes belong to diverse taxa, and the bias of their codon usage is in the range most frequently observed in the genomes of the CUTG database. In all cases, adjustment of the proportion of two compartments in the model within the range 0.55 ≤α ≤0.8 allows the model to reproduce these frequency sequences quite well. Although the theoretical frequency sequences are determined by varying substantial contributions from both hypothesized compartments in these cases, the observed sequences are reproduced without any change made to the distributions that are assumed to underly codon usage in each compartment.

**Figure 4:**
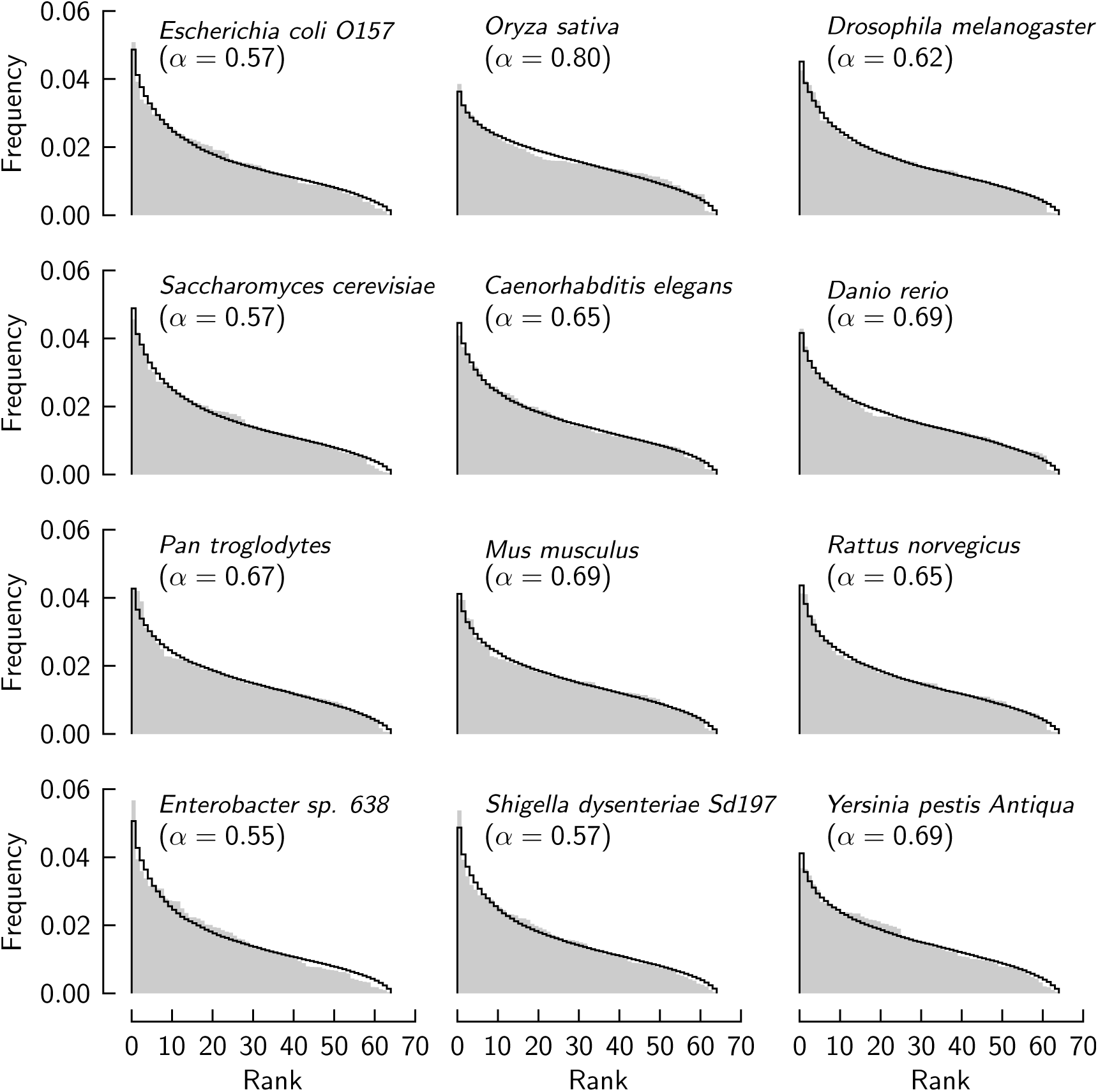
Observed frequencies of codon occurrence compared to theory. Sequences of frequencies of 12 genomes (*gray-shaded*). *Lines* are computed from theory with the indicated proportion, *α*, of the compartment with Gaussian distribution.

Fig. 5A,B summarize the application of the model to the entire set of genomes, divided between eubacteria and archaea (A) and plants, invertebrates, and vertebrates (B). The proportion of the Gaussian compartment, *α*, which is obtained by the fits of the model to the data, is plotted versus the observed bias (eq. 1) for each individual genome (*symbols*). The *lines* show the theoretical relation

**Figure 5:**
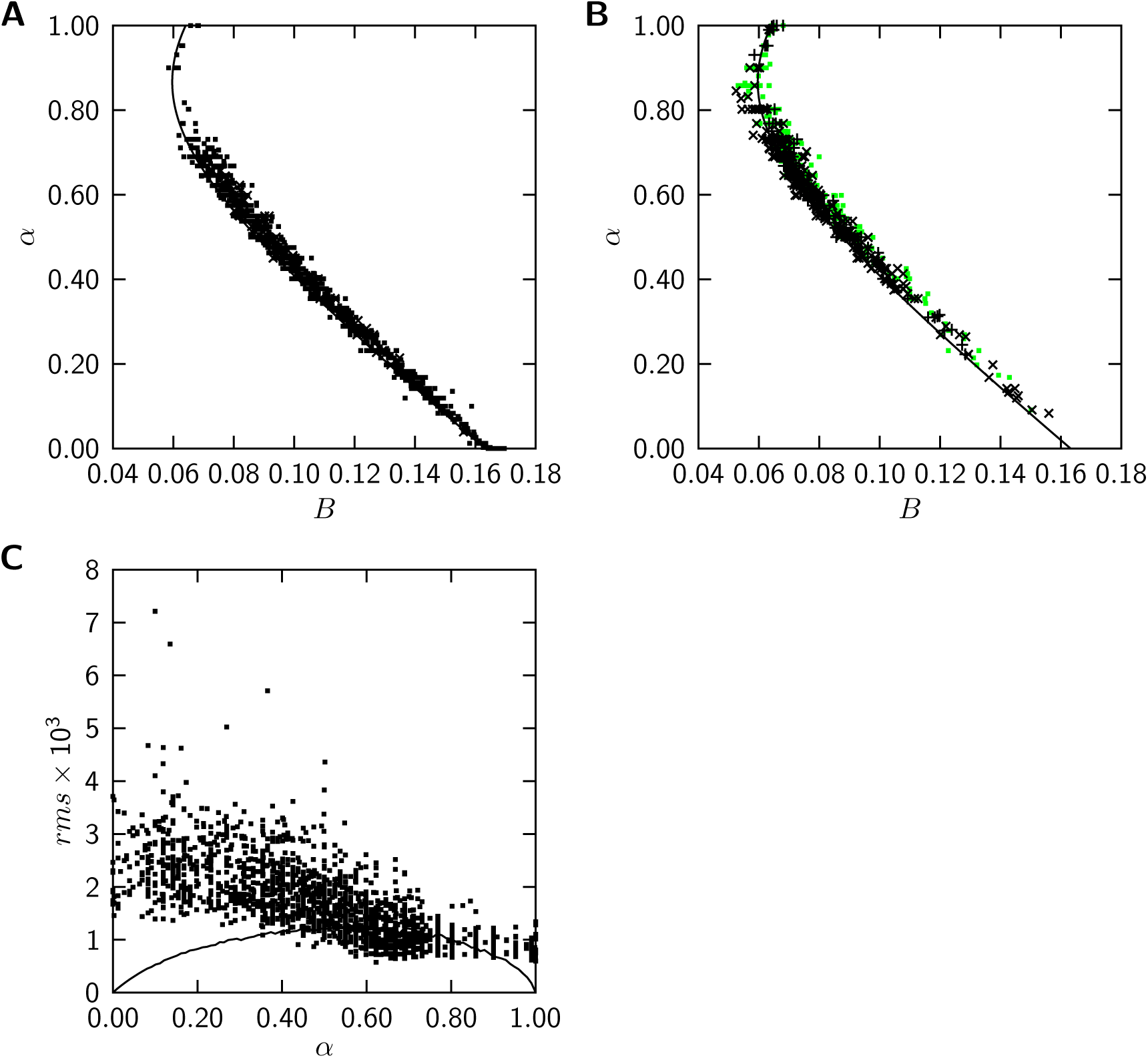
The two-compartment model describes biased codon usage of 1840 genomes. (A) The single varied model parameter *α* describing a genome plotted versus the observed codon usage bias, eq.1:*black symbols* eubacteria, *red symbols* archaea. The *line* represents the theoretical relation, eq. 6.(B) Like panel A, representing eukaryotes: *green* plants, *blue* invertebrates, *red* vertebrates. (C) rms of residuals between the observed and theoretical frequency/rank relations (*symbols*) and theoretical rms expected to arise due to unknown ranks of a different codon in the model compartments (*line*).

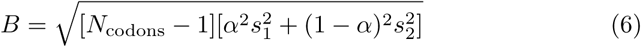

where *s*_1_ = 0.0081 is the standard error of the truncated Gaussian distribution, and *s*_2_ = 0.0207 the standard error of the empirical distribution.

Genomes follow the theoretical relation very closely for all taxa represented here, showing that the model produces a consistent relation between compartment proportions and codon usage bias over the full range of bias and of shapes found in the frequency/rank relations of these 1840 genomes.

The accuracy of the model in describing these data is quantified in Fig. 5C showing the rms residual between observed and theoretical frequency/rank relations for each genome (*symbols*). The *line* gives the rms residual expected to arise from the uncertain frequency ranks of a different codon in the two model compartments (see Construction of the model and Fig. 2E). For the majority of genomes, the actual residual of the fit is similar to the residual expected from the uncertainty of ranks.

## 5. Discussion

We have analyzed aspects of biased usage of different codons at a scale ranging from the individual full genome to many genomes of different taxa. Building on work of Gussein-Zade and Borodovsky [19], we have constructed a statistical model that describes the ordered sequence of codon frequencies of a genome, and the variations of the ordered frequency sequence across genomes.

The model posits that codon usage is generally inhomogeneous within a genome, albeit to varying extents. Two compartments of codons, each with a distinct distribution of codon frequencies, provide a good description if the proportion of the two compartments is adjusted for the specific genome while the compartmental frequency distributions are kept the same. The model parameter distinguishing the codon usages of different genomes thus has a simple meaning.

The two distributions of the model are ‘antipodal’ in that one is Gaussian with a scale that results in weak bias of codon usage, and the other, described empirically, results in very strong bias. The model is not concerned with a specific map relating the two compartments to the genome’s codons (at whatever scale), nor with a specific association of different codons (or their nucleotide composition) with rank in the ordered frequency sequence. Nevertheless, the model reveals a pattern of codon usages that is apparently universal among genomes.

The existence, within one and the same genome, of two forms of codon usage indicates the existence of ‘inhomogeneous conditions of molecular evolution within a genome’ [19]. Inhomogeneity must be strong enough to drive codon usage into into two antipodal compartmental patterns. Of the conditions that support either antipodal distribution of codon frequency, the ones that result in the strongly biased distribution might intuitively appear as the stronger determinants of codon usage. Then, the compartment with Gaussian distribution might be determined more by the absence of strong determinants than by distinguished conditions that favor the more balanced codon usage. Such a simple view of conditions determining codon usage is diαracted by the fact that even the postulated ‘strong’ conditions must generally result in much different ranks of particular different codons in similar ordered sequences of frequency (Fig. 1B), a tendency recognizable even in genomes that use codons with less bias (Fig. 1A). It is therefore clear that a further discussion of conditions that shape codon usage needs to consider the fate of the specific different codons (a study of this kind will be presented in a separate paper).

In the codon compartment that we model by a Gaussian distribution of frequency, that distribution likely summarizes consequences of many evolutionary conditions as well as outcomes in many subpopulations of the compartment. The Central Limit Theorem of statistics would then predict that the joint distribution approaches a Gaussian, even if individual contributions themselves do not produce a Gaussian distribution. We find that one Gaussian distribution (of fixed scale and position) describes the compartment across genomes. Conditions creating this distribution might be universal across genomes but must have weak consequences individually. On the other hand, the strongly biased distribution of the other compartment, by virtue of being non-Gaussian, suggests a comparably simple structure informed by few but strong evolutionary conditions. This raises the question of how consequences of both weak and strong evolutionary conditions might co-exist in one genome. An answer might have to include not only the short-term evolutionary status but also the long-term evolutionary history of an organism – the evolutionary conditions informing codon compartments need not co-exist within one and the same span of time.

## 6. Acknowledgments

Research reported in this publication was supported by the National Institute on Aging of the National Institutes of Health under Award Number T32 AG0047126 (to BK). The content is solely the responsibility of the author(s) and does not necessarily represent the official views of the National Institutes of Health.

